# Human microglia in brain assembloids display region-specific diversity and respond to hyperexcitable neurons carrying *SCN2A* mutation: Microglial diversity and response in assembloids

**DOI:** 10.1101/2025.06.04.657874

**Authors:** Jiaxiang Wu, Xiaoling Chen, Jingliang Zhang, Kyle Wettschurack, Morgan Robinson, Weihao Li, Yuanrui Zhao, Ye-Eun Yoo, Brody A. Deming, Akila D. Abeyaratna, Zhefu Que, Dongshu Du, Matthew Tegtmeyer, Chongli Yuan, William C. Skarnes, Jean-Christophe Rochet, Long-Jun Wu, Yang Yang

**Affiliations:** Borch Department of Medicinal Chemistry and Molecular Pharmacology, College of Pharmacy, Purdue University, West Lafayette, IN 47907, USA; Purdue Institute for Integrative Neuroscience, Purdue University, West Lafayette, IN 47907, USA; Davidson School of Chemical Engineering, College of Engineering, Purdue University, West Lafayette, IN 47907, USA; ENT Institute and Department of Otorhinolaryngology, Eye & ENT Hospital, Fudan University, Shanghai 200031, China; School of Life Sciences, Shanghai University, Shanghai 200444, China; Department of Biological Sciences, Purdue University, West Lafayette, IN 47907, USA; The Jackson Laboratory for Genomic Medicine, Farmington, CT 06032, USA; Center for Neuroimmunology and Glial Biology, Institute of Molecular Medicine, University of Texas Health Science Center at Houston, Houston, TX 77030, USA

**Keywords:** microglia, brain organoid, assembloid, midbrain-striatal circuit, Na_V_1.2

## Abstract

Microglia critically shape neuronal circuit development and function, yet their region-specific properties and roles in distinct circuits of the human brain remain poorly understood. In this study, we generated region-specific brain organoids (cortical, striatal, and midbrain), each integrated with human microglia, to fill this critical gap. Single-cell RNA sequencing uncovered six distinct microglial subtypes exhibiting unique regional signatures, including a subtype highly enriched for the GABA_B_ receptor gene within striatal organoids. To investigate the contributions of microglia to neural circuitry, we created microglia-incorporated midbrain-striatal assembloids, modeling a core circuit node for many neuropsychiatric disorders including autism. Using chemogenetics to activate this midbrain-striatal circuit, we observed increased calcium signaling in microglia involving GABA_B_ receptors. Leveraging this model, we examined microglial responses within neural circuits harboring an *SCN2A* nonsense (C959X) mutation associated with profound autism. Remarkably, microglia displayed heightened calcium responses to *SCN2A* mutation-mediated neuronal hyperactivity, and engaged in excessive synaptic pruning. These pathological effects were reversed by pharmacological inhibition of microglial GABA_B_ receptors. Collectively, our findings establish an advanced platform to dissect human neuroimmune interactions in sub-cortical regions, highlighting the important role of microglia in shaping critical circuitry related to neuropsychiatric disorders.

**Graphical abstract:** 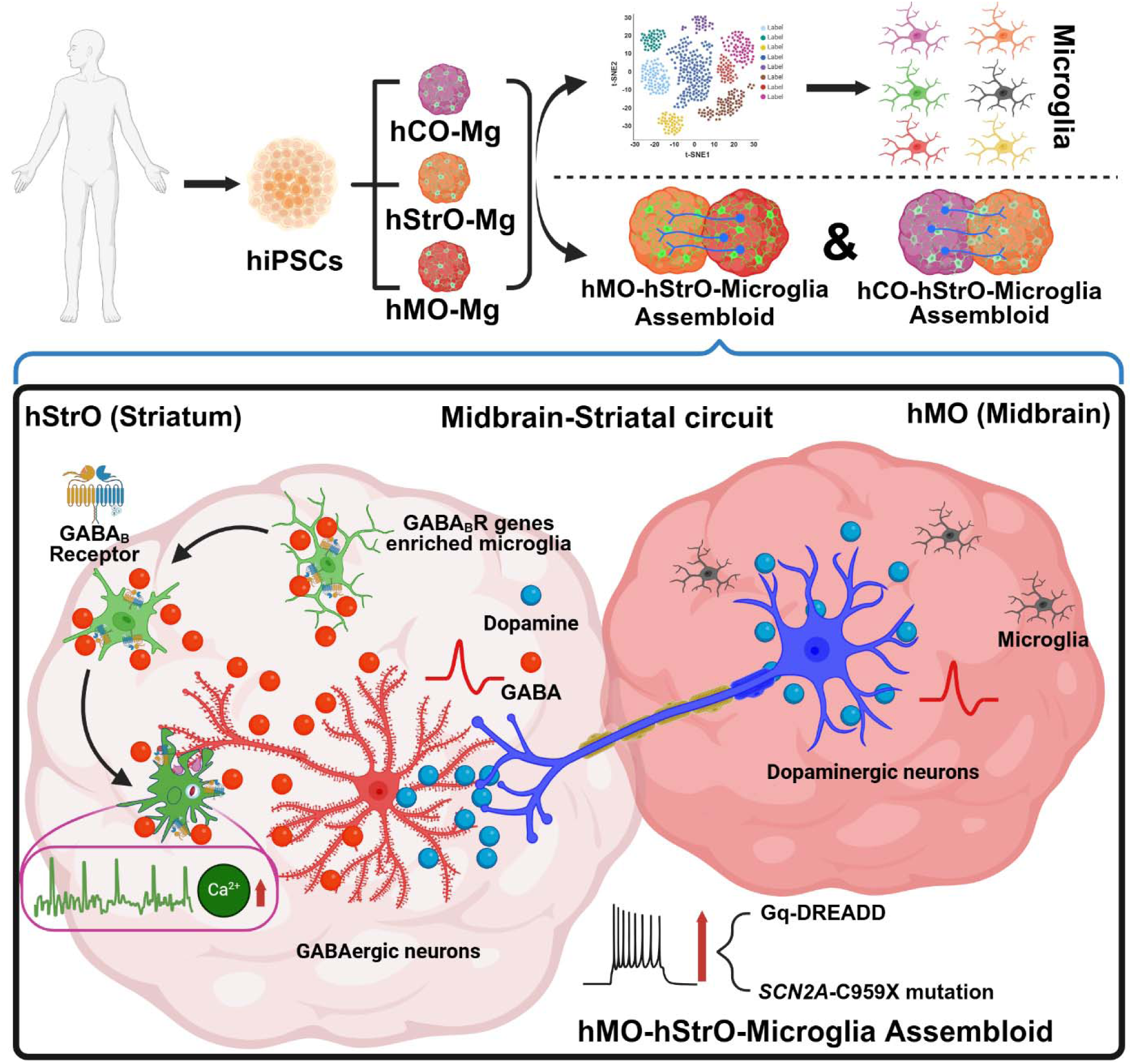

**Teaser:** Modeling regional microglial diversity in sub-cortical regions is challenging. We generated human organoid and assembloid models containing microglia that acquire region-specific heterogeneity. Our work shows dynamic responses of microglia when exposed to hyperexcitable midbrain-striatal circuits, providing an exciting platform to study neuroimmune interactions in human brain development and neuropsychiatric disorders, including *SCN2A* mutation-mediated monogenic autism.

**Highlights:**

• Single-cell RNA sequencing analyses reveal six distinct microglial subtypes that spontaneously attain unique specialization in human cortical, striatal, and midbrain organoids.
• Microglia facilitate axonal projections across regional organoids, promoting assembloid formation.
• Microglia respond to hyperexcitable neurons via calcium signaling and exhibit excessive pruning of neuronal synapses.
• Blocking microglial GABA_B_ receptors normalizes calcium activity and reduces synaptic pruning, suggesting a potential targeting strategy for synaptic deficits.

## INTRODUCTION

Microglia, the resident immune cells of the brain, originate from erythromyeloid progenitors in the yolk sac and migrate into the developing brain during early embryogenesis, subsequently differentiating and maturing(*1, 2*). Beyond their classical functions as immune cells, microglia play indispensable roles in regulating neuronal maturation, synaptic refinement, and the assembly of cross-regional neural circuits throughout development(*3–7*). Notably, emerging evidence from rodent studies reveals remarkable region-specific identities of microglia that display spatial heterogeneity and function distinctly in different neuronal circuits and microenvironments(*8–11*). Interestingly, bulk and single-cell transcriptomic analyses further reveal gene-expression profile-defined subpopulations of microglia across brain regions and developmental stages in rodent models(*12, 13*). Despite these advancements, the regional heterogeneity of human microglia particularly in sub-cortical structures, such as the striatum and midbrain highly relevant to human neuropsychiatric disorders(*14, 15*), remains underexplored.

Indeed, microglial dysfunction has been strongly implicated in numerous neuropsychiatric disorders, including depression, schizophrenia, and autism spectrum disorder (ASD)(*16–18*). In our recent study, we identified microglial abnormalities in a monogenic form of ASD caused by a nonsense mutation in *SCN2A*, which encodes the voltage-gated sodium channel Na_V_1.2(*16*). Using an ASD-associated *Scn2a*-deficient mouse model, we further observed aberrant microglial-mediated synaptic pruning, alongside deficits in synaptic formation and neuronal hyperexcitability(*16, 19, 20*). Interestingly, microglia were also found to respond to hyperexcitable neurons through alterations in their calcium signaling(*21, 22*), which is known to be closely correlated with their phagocytic activity and synaptic pruning function(*23*). However, how human microglia behave within the critical midbrain-striatal circuits related to ASD remains unclear.

To address these gaps, here we developed human induced pluripotent stem cell (hiPSC)-derived organoid and assembloid models integrated with microglia, allowing detailed exploration of microglial heterogeneity and neuroimmune interactions in human cell-based models. Single-cell RNA sequencing analysis identified distinct region-specific microglial subtypes, with striatal microglia notably enriched for genes encoding GABA receptors, notably GABA_B_ receptors. Using chemogenetics and live calcium imaging, we observed that striatal microglia actively respond to the activation of the midbrain-striatal circuits involving GABA_B_ receptors. Leveraging this advanced model, we revealed that the microglia can respond to hyperexcitability caused by an ASD-linked *SCN2A* nonsense mutation, exhibiting elevated calcium signaling and excessive synaptic pruning. Moreover, we found that pharmacological inhibition of GABA_B_ receptors restored microglial activity to baseline and mitigated inhibitory synapse loss, highlighting a novel GABA_B_ receptor-dependent axis underlying neural circuit dysfunction in ASD.

## RESULTS

### hCO-Mg, hStrO-Mg, and hMO-Mg models reveal spatially correlated microglial heterogeneity in distinct human brain organoids

To investigate the spatiotemporal characteristics of human microglia in distinct brain regions composed of different neuronal types, we generated region-specific neural organoids alongside unspecialized/naïve microglia derived from hiPSCs (**Figure S1A**). These unspecialized microglia were subsequently co-cultured with cortical, striatal, or midbrain organoids, establishing three distinct models: human cortical organoids with microglia (hCO-Mg), human striatal organoids with microglia (hStrO-Mg), and human midbrain organoids with microglia (hMO-Mg) (**Figure 1A**). These co-culture models enable microglial infiltration into specific brain regions, allowing microglia to naturally adopt regional properties from their local neural environment. Initially, microglia localized to organoid peripheries but achieved complete infiltration and uniform distribution within 8 weeks of co-culture (**Figure 1B, C**). Regional identities of organoids were verified by specific neuronal markers: T-box brain protein 1 (TBR1) for hCO-Mg, GABA for hStrO-Mg, and tyrosine hydroxylase (TH) for hMO-Mg (**Figure 1D**). Microglial morphology changed over time, transitioning from larger soma with fewer processes to smaller soma with complex branching, which represents a more mature stage (**Figure S2A–C**). Our results suggest successful microglial integration into organoids.

**Figure 1.**
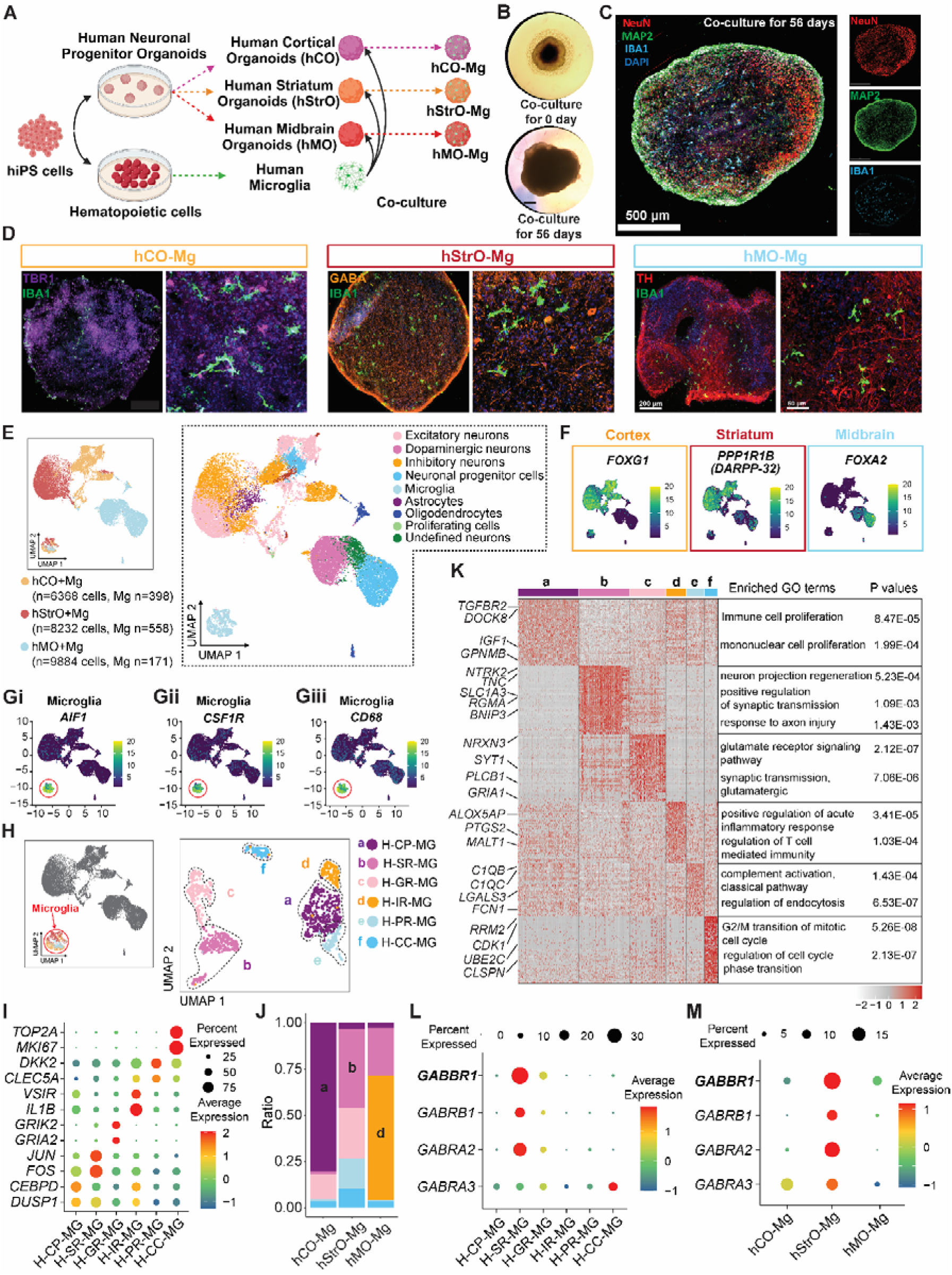
hCO-Mg, hStrO-Mg, and hMO-Mg models reveal spatially distinct microglial heterogeneity within different human brain organoids. (A) Schematic describing the generation of 3D human cortical organoids incorporating microglia (hCO-Mg), human striatal organoids incorporating microglia (hStrO-Mg), and human midbrain organoids incorporating microglia (hMO-Mg) from hiPSCs. (B) Representative brightfield images of striatal organoids co- culture with microglia for 0 days and 56 days. Scale bar: 220 μm. (C) Representative quadruple- staining images (NeuN, Map2, IBA1, and DAPI) from striatal organoids co-culture with microglia for 56 days. Scale bar: 550 μm. (D) Representative images of hCO-Mg (TBR1 and IBA1), hStrO-Mg (GABA and IBA1), and hMO-Mg (TH and IBA1). Full-size images (0.5x) scale bar: 200 μm, zoomed-in images (20x) scale bar: 50 μm. (E) Uniform Manifold Approximation and Projection (UMAP) plot of integrated hCO-Mg, hStrO-Mg, and hMO-Mg. (F and G) UMAP visualization of cerebral cortical marker: *FOXG1*, striatal marker: *PPP1R1B*, midbrain marker: *FOXA2*, and (Gi-Giii) microglial markers: *AIF1*, *CSF1R*, and *CD68*. (H) UMAP visualization of the microglia with 6 clusters. H: human, MG: microglia, CP: cell proliferation, SR: stimulus- response, GR: glutamate-related response, IR: immune response, PR: pruning response, CC: cell cycle. (I) The Dot plot shows the percentage of selected markers for each microglial cluster. (J) The distribution of microglial clusters in different brain region organoids model. (K) Heatmap displaying the top 50 genes of each microglial cluster. Distinct genes (left) related to major types are highlighted with enriched Gene Ontology terms (right). (L and M) Dot plot shows the percentage and average expression of GABA receptor genes in (L) different microglial clusters and (M) microglia from different brain region organoids.

To understand the molecular profiles of microglia within these organoids, we conducted single-cell RNA sequencing (scRNA-seq). Transcriptomic analysis of integrated datasets identified nine clusters using Uniform Manifold Approximation and Projection (UMAP). The highly expressed region-specific neuronal marker genes (*FOXG1, CAMK2A, GRIA2* for hCO; *PPP1R1B, DLX1, GAD1, GAD2* for hStrO; and *FOXA2, SHH* for hMO) validated regional organoid identities (**Figures 1E, F**; **S1B–I**). Microglial marker genes (*AIF1*, *CSF1R*, *CD68*) distinctly identified microglia clusters on the UMAP plot (**Figure 1G**). Further sub-clustering revealed six distinct microglial subpopulations (clusters a-f) (**Figure 1H, I**): cluster a (H-CP-MG) is marked by *DUSP1*, *CEBPD*(*24, 25*); cluster b (H-SR-MG) expresses stimulus-responsive genes *JUN*, *FOS*; cluster c (H-GR-MG) is enriched in glutamate-related genes *GRIK2*, *GRIA2*; cluster d (H-IR-MG) is featured by immune-response genes *VSIR*, *IL1B*; cluster e (H-PR-MG) expresses synaptic pruning genes *DKK2*, *C1QB*, *FCN1*; and cluster f (H-CC-MG) represents early developmental microglia expressing *TOP2A*, *MKI67* (**Figure 1K**).

To further assess regional microglial heterogeneity, we examined the distribution of microglial subpopulations across organoid types. Results showed striking enrichment patterns (**Figure 1J, K**): Cluster a was predominantly found in hCO-Mg (>75%) and exhibited high expression of *TGFBR2*, *DOCK8*, *IGF1*, and *GPNMB*. These genes were associated with enhanced proliferative capabilities by Gene Ontology (GO) analysis. Cluster b was significantly enriched in hStrO-Mg (∼50%) and demonstrated enriched expression of *NTRK2*, *SLC1A3*, and *BNIP3*. These genes were implicated in the modulation of neuronal excitability and synaptic plasticity by GO analysis. Cluster d primarily populated hMO-Mg (∼70%) and prominently expressed *ALOX5AP*, *PTGS2*, and *MALT1*, indicative of specialized immune-regulatory functions. Furthermore, consistent with prior findings implicating GABA signaling in microglial activity and inhibitory circuit formation(*26*), hStrO-Mg microglia showed an overall enriched expression pattern of multiple GABA receptor-related genes (*GABBR1*, *GABRB1*, *GABRA2*) (**Figures 1M; S2D–F**). In particular, cluster b (H-SR-MG) exhibited predominant expression of these GABA receptor-related genes (**Figure 1L**), correlating with the abundant GABAergic neurons in hStrO and suggesting functional interplay between these microglia and GABAergic neuronal populations. Collectively, our findings provide compelling evidence of region-specific microglial heterogeneity at the transcriptional level within cortical, striatal, and midbrain human organoids, reflecting spatially correlated microglial functional diversity.

### Microglia facilitate axonal projections and enhance assembloid formation

Rodent studies have demonstrated the critical role of microglia in shaping neural connectivity(*27*), influencing both local neural circuits(*9, 26*) and inter-regional projections(*28*). However, the function of microglia in human cell-based neural circuit models remains largely unexplored. Given that the striatum primarily receives projections from cortical glutamatergic neurons and midbrain dopaminergic neurons, we developed assembloid models to reconstruct these circuits by fusing human striatal organoids with microglia (hStrO-Mg) to either human cortical organoids (hCO-Mg) or human midbrain organoids (hMO-Mg) (**Figure 2A**). After generating cortical-striatal-microglia assembloids (hCO-hStrO-Mg) and midbrain-striatal-microglia assembloids (hMO-hStrO-Mg), regional specificity was confirmed using marker expressions: TBR1 and CTIP2 for cortical organoids, GAD67 and DRD1 for striatal organoids, and FOXA2 and OTX2 for midbrain organoids (**Figure 2B**). Furthermore, we validated that each organoid maintain the presence of microglia after fusion by performing co-immunostaining for GABA/IBA1 in cortical-striatal assembloids and TH/IBA1 in midbrain-striatal assembloids, confirming integration of neuron and microglia (**Figure 2C, D**).

**Figure 2.**
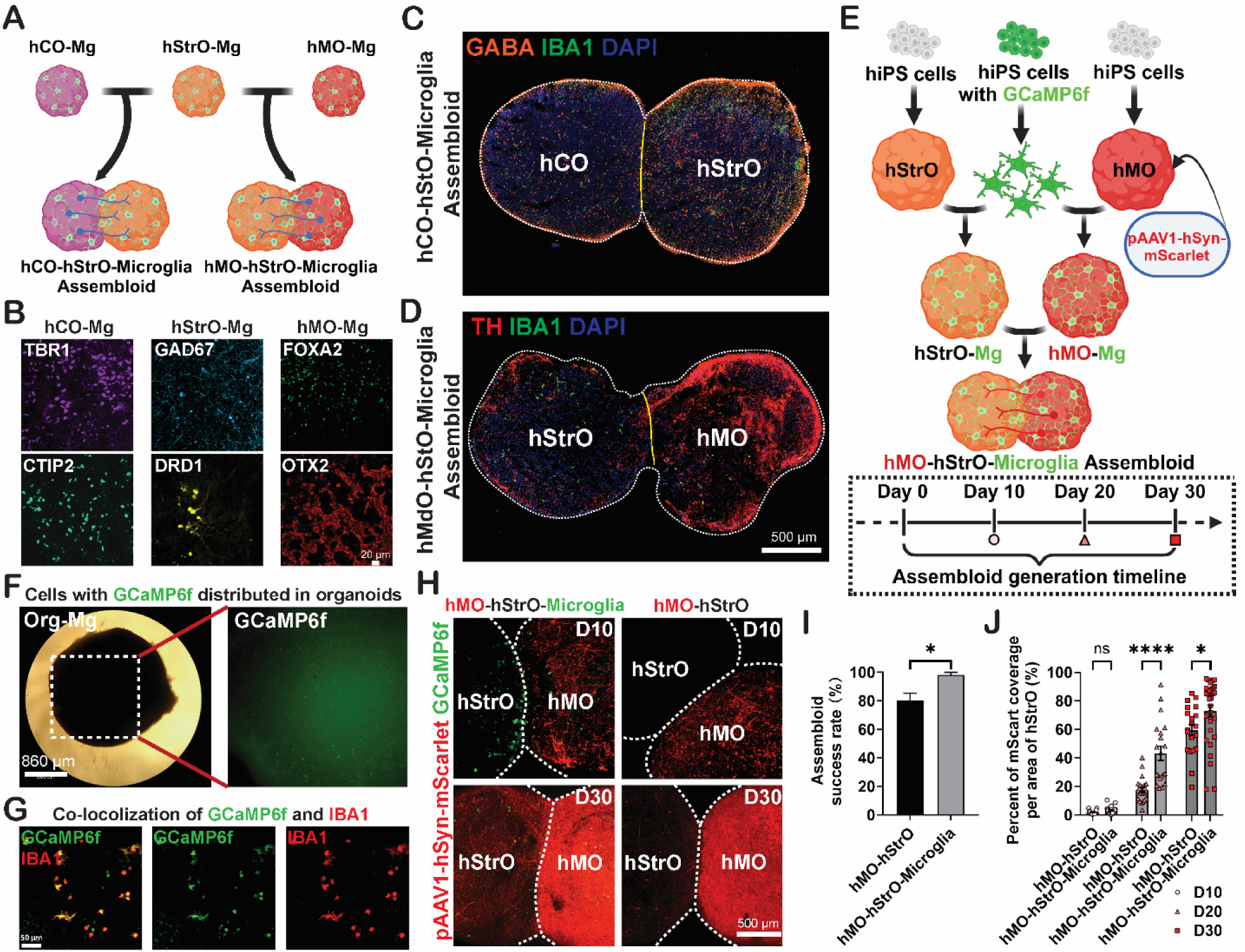
Microglia facilitate axonal projections across distinct brain regions and promote the formation of assembloids. (**A**) Schematic illustrating the microglia-integrated cortico-striatal pathway and midbrain striatal pathway in assembloid models. (**B**) Immunostaining for specific regional markers, hCO-Mg: TBR1 and CTIP2, hStrO-Mg: GAD67 and DRD1, hMO-Mg: FOXA2 and OTX2. Scale bar: 20 μm. (**C** and **D**) Representative image of 3D (C) hCO-hStrO-Microglia assembloid and (**D**) hMO-hStrO-Microglia assembloid. Scale bar: 500 μm. (**E**) Schematic illustrating the hMO-hStrO-Microglia assembloid by using GCaMP6f hiPS cell line to generate microglia and AAV1 for anterograde viral tracing (top). Schematic describing the assembloid generation timeline (bottom). (**F**) Representative images demonstrating strong GCaMP6f signals in the organoid-microglia models. Scale bar: 860 μm. (**G**) Representative images illustrating microglia carry the GCaMP6f signal in the organoid-microglia models. Scale bar: 50 μm. (**H**) Representative images showing the midbrain striatal projections and presence of GCaMP6f^+^ cells of different groups at day 10 and day 30. Scale bar: 500 μm. (**I**) Quantification of success rate of assembloid formation across different groups. Unpaired Student’s t-test, n=5 batches. *p* < 0.05 (*). (**J**) Quantification of mScart coverage per area of hStrO in different groups. Unpaired Student’s t-test, hMO-hStrO and hMO-hStrO-Microglia at day 10: n=20 assembloids. hMO-hStrO and hMO-hStrO-Microglia at day 20: n=20 assembloids. hMO-hStrO at day 30: n=20 assembloids, and hMO-hStrO-Microglia at day 30: n=30 assembloids. *p* < 0.05 (*), *p* < 0.0001 (****), ns: no significance. Data are represented as mean ± SEM.

The projection of midbrain dopaminergic neurons to striatal neurons forms a classical circuit implicated in various neuropsychiatric disorders(*14*). Yet, the contribution of microglia to this circuitry remains unclear. To elucidate it, we reconstructed midbrain-striatal assembloids (hMO-hStrO-Mg) (**Figure 2E, F**), utilizing microglia derived from genetically modified hiPSC lines expressing the calcium sensor GCaMP6f (engineered at the AAVS safe locus). The use of GCaMP6f-labeled microglia enables the live tracking of microglia and microglial calcium dynamics within these assembloids. Prior to fusion, midbrain organoids were labeled with pAAV-hSyn-mScarlet to visualize midbrain compartments distinctly. Immunostaining confirmed GCaMP6f-labeled microglia successfully integrated into assembloids, indicated by co- localization of IBA1 and GCaMP6f signals (**Figure 2G**).

Live imaging post-fusion revealed a progressive increase in mScarlet-labeled midbrain-to-striatal axonal projections over time (**Figure 2H, J**). Notably, microglia-integrated assembloids showed an initial clustering of microglia at the fusion interface by day 10, with subsequent redistribution at later stages. Moreover, assembloids containing microglia exhibited significantly enhanced mScarlet-labeled axonal projections compared to those without microglia (**Figure 2H, J**), along with notably higher fusion success rates (**Figure 2I**). Together, these findings from our microglia-integrated assembloid model indicate that microglia actively facilitate axonal projections, significantly enhancing the assembly, and thereby promoting functional integration of midbrain-striatal circuits.

### Microglial respond to elevated neuronal activity in the hStrO region of hMO-hStrO assembloids

Microglia are known to respond dynamically to changes in neuronal activity(*29, 30*). Before we investigated how microglia sense neuronal activity within human midbrain-striatal circuits, we first performed experiments to validate the functional connectivity of the hMO-hStrO assembloids. We employed a chemogenetics approach by transducing hMO-Mg neurons with pAAV-hSyn-hM3(Gq)-mCherry, enabling Gq-DREADD-mediated activation of the hMO neurons, which send axonal projections to the hStrO. These transduced hMO-Mg were then fused with hStrO-Mg to generate assembloids (**Figure 3A**). Using high-density multi-electrode arrays (HD-MEA), we recorded neural activity following the application of 10 μM clozapine N-oxide (CNO) to activate Gq-DREADDs in hMO. As expected, CNO robustly increased neuronal firing in the hMO region directly (**Figure 3C left**, **3D**). Notably, this elevated activity was efficiently transmitted to the hStrO compartment, as shown by increased neuronal firing in hStrO as well, confirming functional midbrain-striatal connectivity within assembloids (**Figure 3C right, 3D**).

**Figure 3.**
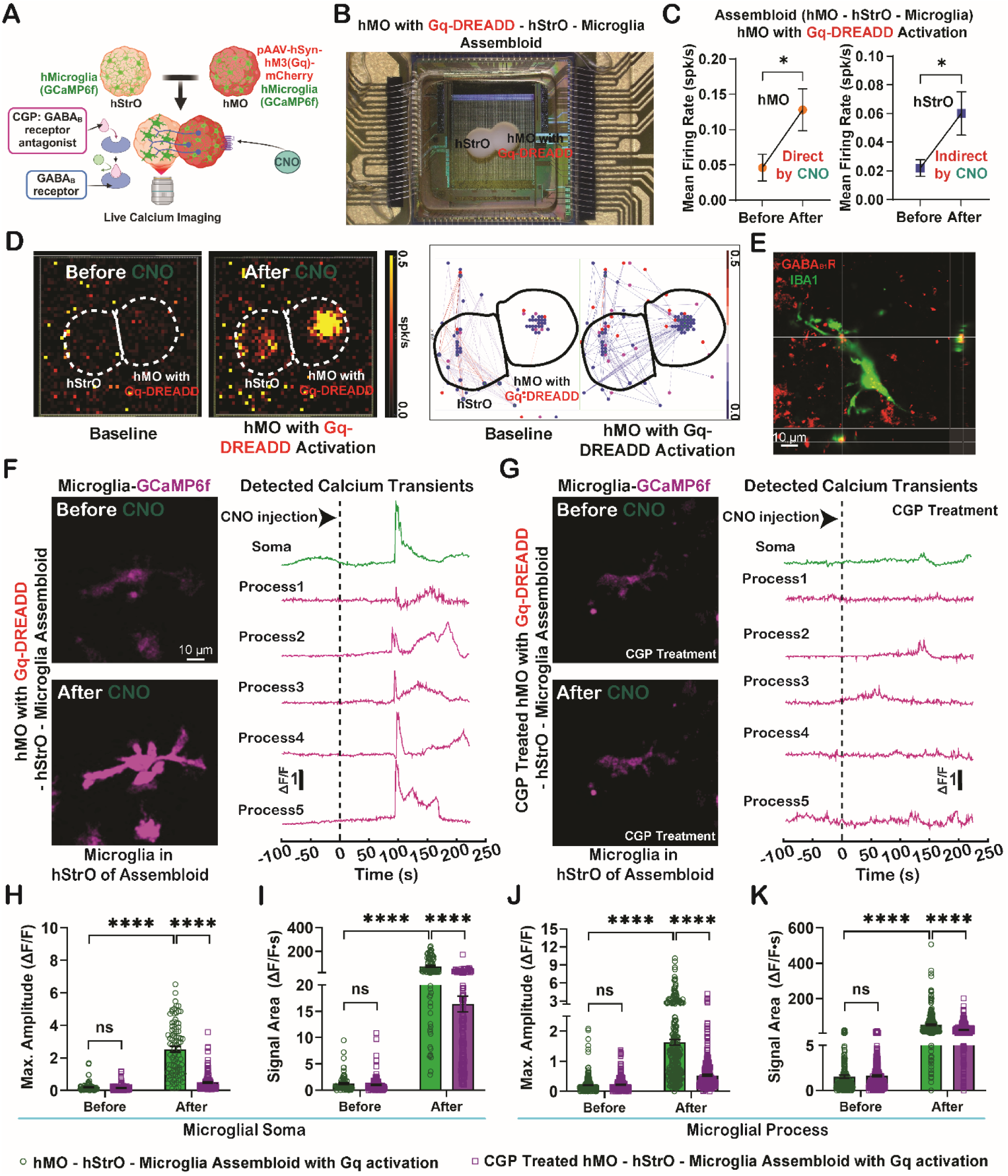
Antagonizing the GABA_B_ receptors by CGP attenuates the microglial responses to elevated neuronal activity in the hStrO of the assembloids. (**A**) Schematic illustrating live calcium imaging of microglia in the hStrO region after CNO exposure. (**B**) Representative image of a hMO-hStrO-Microglia assembloid on a 3-brain high-density microelectrode array (HD-MEA) system. The hMO region was transduced with Gq-DREADD. (**C**) Quantification of mean firing rate in hMO region and hStrO region from the hMO with Gq-DREADD-hStrO-Microglia Assembloid before and after Gq-DREADD activation (CNO exposure). Unpaired Student’s t-test, n=5 assembloids. *p* < 0.05 (*). (**D**) Representative heat map (left) and representative neuronal connection map (right) by HD-MEA recording in the hMO with Gq-DREADD-hStrO-Microglia Assembloid before and after Gq-DREADD activation (CNO exposure). (**E**) Representative orthogonal image of microglia in the hStrO region of hMO with Gq-DREADD-hStrO-Microglia Assembloid. Each cross in the three graphs represents the GABBR1 marker labeled microglial GABA_B_ receptors from three angles. Scale bar: 10 μm. (**F**) Microglial calcium activity (GCaMP6f) in the hStrO region of hMO with Gq-DREADD-hStrO-Microglia Assembloid before and after Gq- DREADD activation (CNO exposure) (left) and representative ΔF/F with detected calcium transients from microglial cells (right). Scale bar: 10 μm. (**G**) Microglial calcium activity (GCaMP6f) in the hStrO region of GABA_B_ receptors antagonist CGP treated hMO with Gq- DREADD-hStrO-Microglia Assembloid before and after Gq-DREADD activation (CNO exposure) (left) and representative ΔF/F with detected calcium transients from microglial cells (right). (**H** and **I**) Quantifications of Max. Amplitude (**H**) and Signal Area (**I**) of calcium signaling in microglial soma in different groups before and after Gq-DREADD activation (CNO exposure). Each group: n=89 microglial somas/20 assembloids. (**J** and **K**) Quantifications of Max. Amplitude (**H**) and Signal Area (**I**) of calcium signaling in microglial process in different groups before and after Gq-DREADD activation (CNO exposure). Each group: n=278 microglial processes/20 assembloids. Two-way ANOVA, *p* < 0.0001 (****), ns: no significance.

Neuronal activity can induce microglial calcium dynamics(*23*). Given prior evidence implicating microglial GABA_B_ receptors in regulating calcium activity(*31*), alongside our scRNA-seq data showing the enrichment of *GABBR1*-expressing microglia in the hStrO, we performed immunostaining in the hStrO region for IBA1 and GABA_B1_R (*GABBR1*), a GABA_B_ receptor subunit. Co-localization of IBA1 and GABA_B1_R confirmed the presence of GABA_B_ receptors on microglia (**Figure 3E**). We next examined how GABA_B_ receptors are involved in calcium dynamics in hStrO microglia using the endogenously expressed calcium sensor GCaMP6f within the midbrain-striatal circuit. Under basal conditions, both microglial somas and processes exhibited minimal spontaneous calcium activity in hStrO (**Figure 3F, H–K**). Following CNO-induced circuit activation, we observed a robust increase in calcium signals within microglial soma and processes in the hStrO (**Figure 3F, H–K**). We then applied CGP 55845 hydrochloride, a selective GABA_B_ receptor antagonist(*31*), to block GABA_B_ receptors. Remarkably, CGP treatment substantially suppressed the elevated microglial calcium signals induced by CNO in both soma and processes of microglia in hStrO (**Figure 3G, H–K**). As GABA_B_ receptors are also expressed in neurons, we assessed whether CGP affected neuronal excitability. Although hStrO neurons showed a trend toward increased excitability upon CGP treatment, this change did not reach statistical significance (**Figure S3A–F**), suggesting that our observed effects were largely from microglia. Together, our data demonstrate that blocking GABA_B_ receptors significantly reduces microglial calcium responses induced by enhanced striatal neuronal activity. Our findings align with previous studies showing that microglia can sense neuronal activity via GABA signaling(*31*) and potentially participate in sculpting inhibitory synapses(*26*).

### Microglia respond to hyperexcitable *SCN2A*-C959X assembloids via GABA_B_ receptor- dependent calcium signaling and excessive synaptic pruning

The midbrain-striatal circuit is strongly implicated in ASD(*15*). Our previous work in a mouse model demonstrated that *Scn2a* deficiency disrupts synaptic transmission and triggers microglial activation, leading to increased synaptic engulfment and impaired neuronal plasticity(*16, 32*). However, the role of microglia in human midbrain-striatal assembloids carrying ASD-causing *SCN2A* mutations remains poorly understood. To address this gap, we established the hMO-hStrO-Mg model carrying a heterozygous *SCN2A* nonsense mutation (C959X) (**Figure 4A**), previously identified in patients with profound ASD(*33, 34*). Electrophysiological analysis revealed significantly elevated network activity in the CX assembloids compared with the isogenic wild-type (WT) control (**Figure S4**), consistent with hyperactivity observed in cortico-striatal assembloids with *SCN2A* deficiency(*35*). To further assess neuronal excitability, we performed patch-clamp recordings of medium spiny neurons (MSNs) in the hStrO region. We revealed that MSNs in the CX assembloids exhibited significantly increased current-evoked action potential firing (**Figure 4B, C**), confirming hyperexcitable phenotypes of the midbrain-striatal circuit with *SCN2A* deficiency.

**Figure 4.**
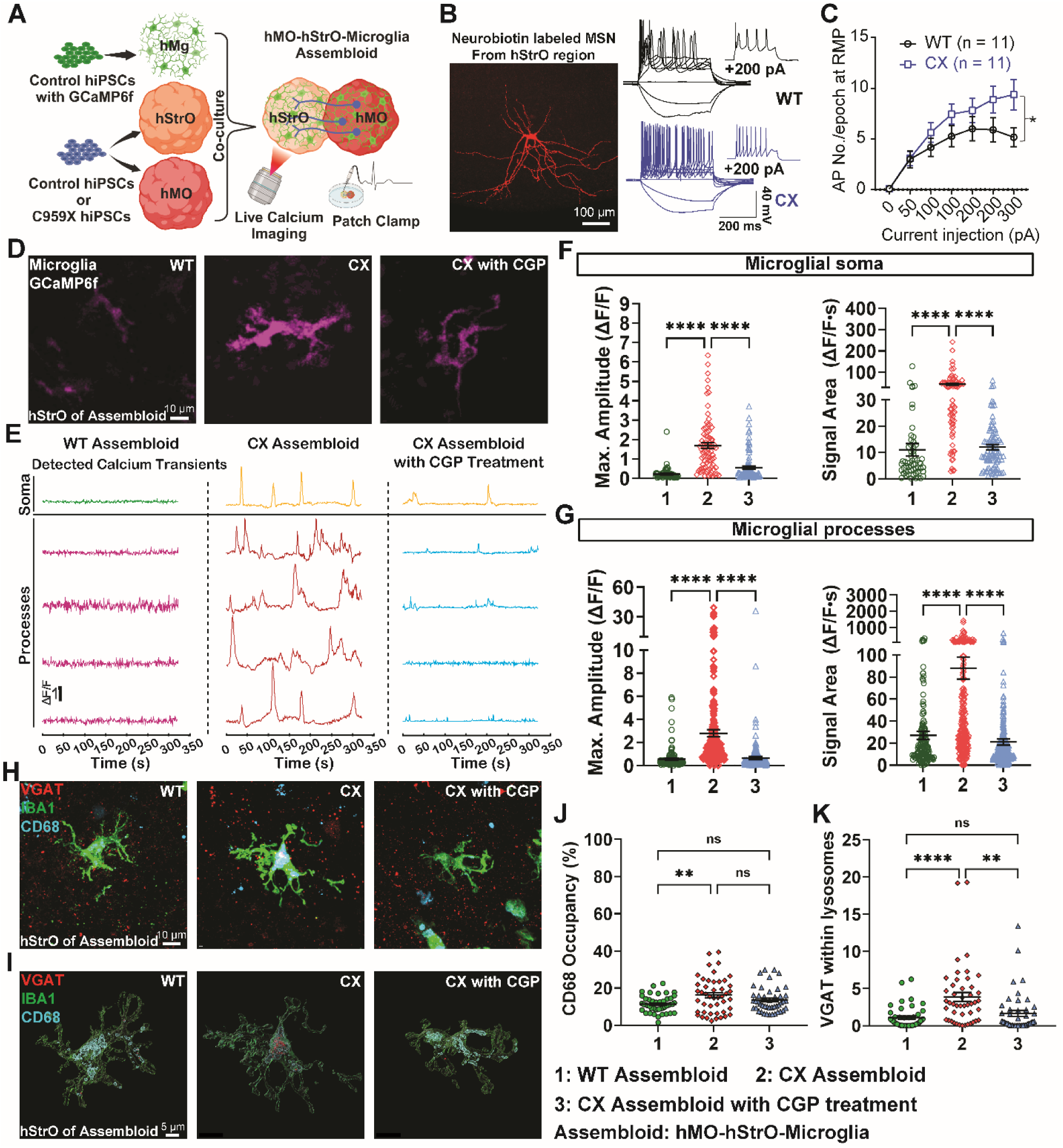
Microglia respond to hyperexcitable assembloids harboring autism-causing *SCN2A* mutation via altered calcium signaling and excessive pruning of synapses. (**A**) Schematic representation of developing microglia-containing *SCN2A* C959X-mutant assembloid. (**B**) A typical medium spiny neuron (MSN) labeled by neurobiotin and representative current-clamp recordings of MSNs from each group. Scale bar: 100 μm. (**C**) The average number of action potentials (APs) generated in response to depolarizing current pulses. Unpaired two-tailed non-parametric Mann-Whitney U test for each current pulse, n=11 from each group. *p* < 0.001 (***). (**D**) Microglial calcium activity (GCaMP6f) in the hStrO region of hMO-hStrO- Microglia Assembloid from each group (WT group, *SCN2A* C959X mutant group, and *SCN2A* C959X mutant with CGP group). (**E**) Representative ΔF/F with detected calcium transients of microglial cells from each group. (**F**) Quantifications of Max. Amplitude and Signal Area of calcium signaling in microglial soma from different groups. WT Assembloid: n=59 microglial somas/15 assembloids, CX Assembloid: n=79 somas/20 assembloids, CX with CGP: n=92 somas/20 assembloids. One-way ANOVA, *p* < 0.0001 (****). (**G**) Quantifications of Max. Amplitude and Signal Area of calcium signaling in microglial process from different groups. WT Assembloid: n=144 microglial processes/15 assembloids, CX Assembloid: n=267 processes/20 assembloids, CX with CGP: n=277 processes/20 assembloids. One-way ANOVA, *p* < 0.0001 (****). (**H** and **I**) Representative triple-staining images (VGAT, IBA1, and CD68) (H) and Imaris reconstructed images from each group (**I**). Scale bar: 10 μm. (**J** and **K**) Quantifications of CD68 positive occupancy (%) (**J**) and VGAT within lysosomes (**K**) from each group. WT Assembloid: n=45 cells/9 assembloids, CX Assembloid: n=46 cells/9 assembloids, CX with CGP: n=45 cells/9 assembloids. One-way ANOVA, *p* < 0.01 (**), and *p* < 0.0001 (****), ns: no significance.

Given the sensitivity of microglial calcium signaling to neuronal activity, we used real-time live-cell calcium imaging to monitor microglial responses in the hStrO region (**Figure 4D, E**). As it was demonstrated that *SCN2A* is a neuron-specific gene that has minimal expression in microglia(*22*), we utilized WT GaMP6f microglia to explore how microglia sense neuronal hyperexcitability in the CX midbrain-stratal assembloids. Compared with the microglia in the WT assembloids, microglia in the CX assembloids displayed significantly elevated calcium activity in both their somas and processes (**Figure 4F, G**). Microglial over-pruning of synapses has been implicated in various neuropsychiatric disorders(*36*). To investigate whether microglia in CX assembloids contribute to synaptic dysfunction in our model, we performed triple immunofluorescence staining for VGAT, IBA1, and CD68. 3D reconstruction and volumetric analysis using Imaris revealed a substantial increase in lysosomal volume (**Figure 4J**) and enhanced phagocytosis of VGAT-positive inhibitory presynaptic terminals by microglia in the CX assembloids (**Figure 4K**), indicating excessive pruning of inhibitory synapses. Moreover, pharmacological inhibition of GABA_B_ receptors using CGP 55845 significantly reduced microglial calcium signaling in the CX assembloids. Notably, this reduction in calcium activity was accompanied by decreased VGAT engulfment (**Figure 4D–G**), which could ultimately alleviate synaptic deficits in *SCN2A*-deficient MSNs. Taken together, our findings demonstrate that microglia could respond to *SCN2A*-C959X-induced neuronal hyperactivity via GABA_B_ receptor-involving calcium signaling, and perform excessive pruning of inhibitory synapses. These results highlight a critical neuroimmune mechanism contributing to synaptic pathology in human cell-based ASD models and suggest a potential strategy targeting microglial GABA signaling for intervention.

## DISCUSSION

Brain organoids and assembloids have rapidly advanced over the past decade, making it possible to study neurodevelopment and neuropsychiatric disorders in 3D structurally organized human cells. While microglia are critical in regulating neuronal function and circuitry(*37*), the region-specific identities of microglia and their role in sub-cortical circuits remain largely unknown. In this study, using scRNA-seq, we revealed distinct, region-specific microglial subtypes, with striatal microglia showing notable enrichment in the GABA_B_ receptor gene. Utilizing chemogenetics combined with live-cell calcium imaging, we demonstrated that these striatal microglia dynamically respond to elevated neuronal activity in midbrain-striatal circuits. Importantly, we found microglia can also respond to hyperexcitable circuitry harboring autism-causing *SCN2A* protein-truncating variant C959X, by increasing microglial calcium signaling, and performing excessive synaptic pruning. Moreover, we found that pharmacological inhibition of GABA_B_ receptors effectively normalized microglial activity and mitigated inhibitory synaptic loss in neurons. Our study thus demonstrates the utility of a powerful platform to understand the roles of microglia in human brain circuitry and neuropsychiatric disorders.

Rodent and post-mortem human tissue studies have revealed that microglia exhibit region-specific phenotypes, including distinct morphologies, cell densities, and gene/protein expression profiles(*38, 39*). Such region-specific features of microglia are thought to be shaped by developmental timing, local neuronal identities, and microenvironmental cues(*8–10, 12, 13, 40, 41*). These dynamic local signals may prompt microglia to perform distinct functions related to synaptic pruning as well as facilitate circuit maturation and axonal projection based on the local context. Given that most insights into microglia diversity to date come from rodent models or post-mortem human tissues, our study fills a critical gap by establishing human brain circuit models to study the identities and functions of human microglia. Using single-cell transcriptomic analysis, we identified six distinct microglial subtypes across three region-specific brain organoids, with each region displaying a characteristic distribution of these subpopulations. Notably, three subtypes (a, b, and f) we identified from our study to a certain degree correspond to microglial populations observed in the early developing human brain tissues(*39*), strengthening the utility of our platform to model human microglia in brain development. In particular, it is worth mentioning that the majority of microglia in hStrO-Mg are type b microglia, showing significant enrichment of GABA_B_ receptor gene expression. This is interesting, considering the abundance of GABAergic neurons in the striatum. Notably, this finding also aligns with evidence from mouse studies, where regional environments (e.g., GABA- or glutamate-enriched circuits) drive transitions in microglial subtypes(*29, 42*). Together, we suggest that our novel human cell-based models provide an advanced platform that, when complemented by animal models and human tissues, can comprehensively elucidate the functions of microglial subtypes across different brain regions and species.

Microglia are known to prune synapses as a way to regulate neuronal network excitability(*29*). In particular, previous studies have found that microglia can dampen neuronal hyperexcitability in chemically triggered seizure mouse models and human cells carrying seizure-related genetic mutations(*22, 30, 43*). Interestingly, it is also found that microglial calcium dynamics are highly related to microglial phagocytosis as well as neuronal excitability monitoring and modulation(*21, 23*). While these results are exciting, they are limited by the existing technologies, which only allow the live-cell imaging of microglia either in 2D culture/brain slices that do not have intact long-range circuitry connectivity, or superficial cortical regions limited by the imaging depth of *in vivo* two-photon microscopy(*21, 44*). Our current platform, therefore, provides a complementary approach to studying human microglia in sub- cortical circuits. Notably, our finding in the midbrain-striatal circuit is consistent with the published results that microglia display elevated calcium signaling in a hyperexcitable environment and perform over-pruning of synapses in disease models of autism/epilepsy(*16, 22*). Moving beyond confirming published results, our study was able to reveal a population of GABA_B_-enriched microglia in hStrO. We further demonstrated that the elevated calcium signal and synaptic over-pruning are microglial GABA_B_ receptor-dependent, providing new insights into the region-specific function of human microglia in regulating neuronal excitability that is not likely to be revealed by existing models.

Our study may have several limitations: **1)** Our assembloid system lacks a vascular component. The brain vasculature, however, is known to maintain homeostasis and influence microglial development and functions(*45, 46*). **2)** While our system can model the classic midbrain to striatum long-range axonal projections, neurons in the striatum should also have long-range axonal projections to various other brain regions(*47–49*), which our current system does not model. **3)** We proposed the GABA_B_ receptors expressed in microglia are important in mediating microglial calcium dynamics and pruning. Using pharmacological blockade of GABA_B_ receptor, we observed greatly diminished Ca^2+^ response in microglia and reduced synaptic over-pruning. Our finding is supported by published results showing microglial GABA_B_ receptors are involved in its phagocytic functions(*26*). However, it is obvious that the GABA_B_ receptors are also expressed in the neurons and pharmacological agents could block GABA_B_ receptors in neurons as well. We reason that neuronal GABA_B_ receptors are unlikely to dominate the observed effects. This is because blocking neuronal GABA_B_ receptors would theoretically enhance neuronal excitability, subsequently increasing microglial sensing of neuronal hyperactivity and elevating calcium signaling. In contrast, our data showed significantly reduced microglial calcium dynamics upon pharmacological blockade of GABA_B_ receptors, supporting a primarily microglia-driven mechanism.

In summary, our study established an advanced platform with substantial potential for investigating microglial functions in neural circuits and their dysfunction in human diseases. It is especially exciting that microglial spatial heterogeneity in the human brain can be partially modeled *in vitro* with human cell-based brain organoids and assembloids. Our findings enable a detailed exploration of potentially diverse microglial roles across different brain regions, including sub-cortical areas that have historically been challenged to access by *in vivo* imaging technology. Moreover, our model allows live-cell imaging and precise experimental manipulations of microglia in a human cell-based system, advancing our understanding of microglial biology and its contribution to human neuropsychiatric diseases.

## ACKNOWLEDGMENTS

We thank Dr. Adam Kimbrough from Purdue University for kindly providing access to Imaris software. The research reported in this publication was partially supported by the NINDS of the NIH (R01NS117585 and R01NS123154 to Y.Y.). X.C. is supported by the American Epilepsy Society (AES) Postdoctoral Research Fellowship. The authors gratefully acknowledge support from the Familie*SCN2A* Foundation for the Hodgkin-Huxley Award to Y.Y. and the Action Potential Grant support to X.C., J.Z., and Y.E.Y. The authors thank all other members of the Yang laboratory at Purdue University for insightful discussions.

## AUTHOR CONTRIBUTIONS

J.W. and Y.Y. conceived and designed the experiments. J.W., X.C., J.Z., K.W., and M.R. performed experiments and analyzed the data. W.C.S. designed and performed the *SCN2A* gene editing experiment. W.L. and Y.Z. participated in data analysis. Y.-E.Y., B.D., A.A., and Z.Q., participated in performing experiments. D.D., M.T., C.Y., W.S., J.-C.R., and L.-J.W. participated in the experimental design. Y.Y. supervised the project. J.W., J.Z., and Y.Y. wrote the manuscript with input from all authors.

## DECLARATION OF GENERATIVE AI AND AI-ASSISTED TECHNOLOGIES IN THE WRITING PROCESS

During the preparation of this work, the authors used ChatGPT 4.5 to improve the readability and language in this manuscript while ensuring that the main conclusions remained unchanged. After using this tool, the authors reviewed and edited the wording as necessary and take full responsibility for the content of the publication.

---

**s-Figure 1.**
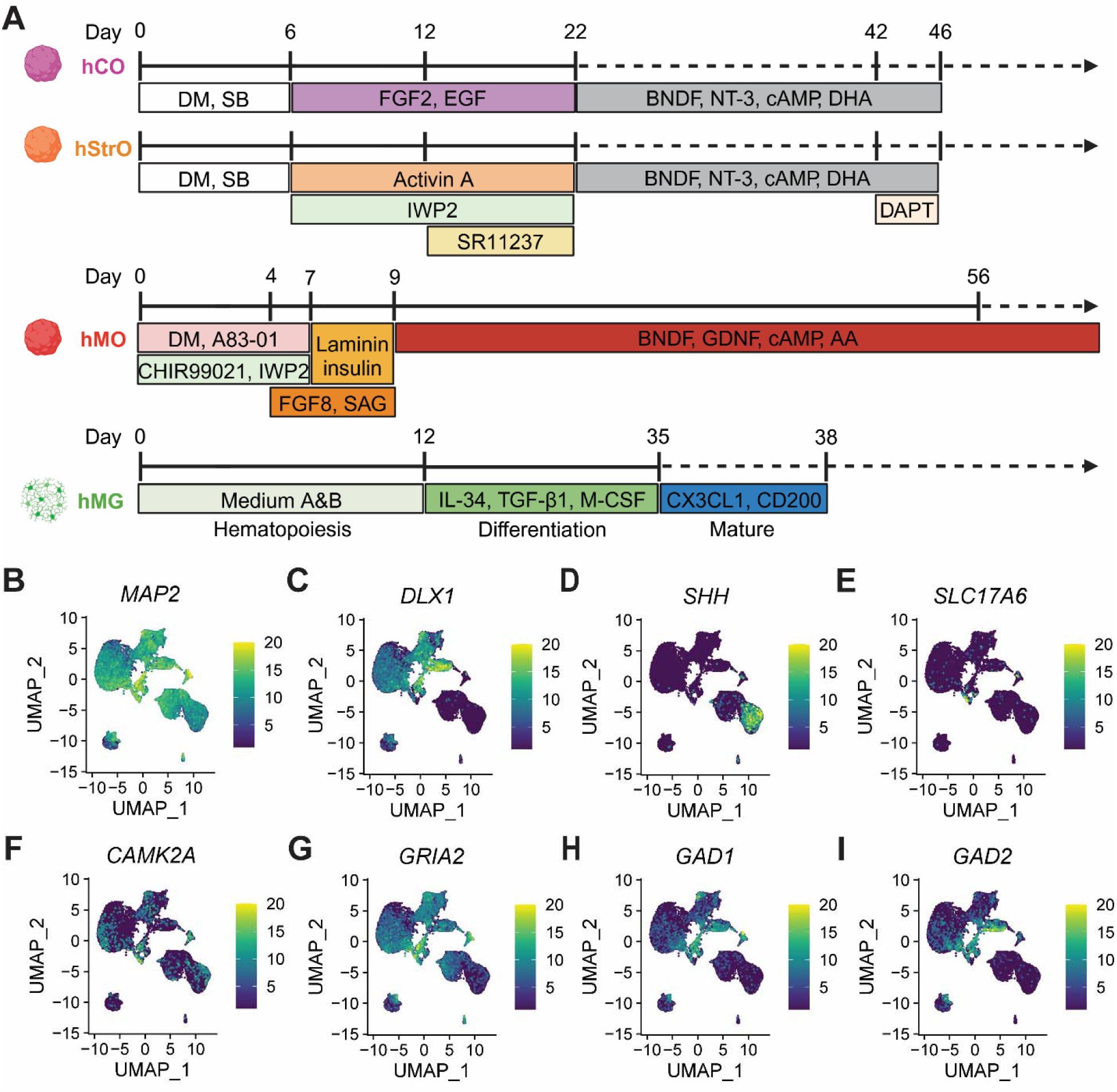
Protocols and single-cell analysis of microglia-integrated brain region- specific organoids. (**A**) Schematic representation of the differentiation protocols for different human brain organoid models. hCO (Human Cortical Organoid), hStrO (Human Striatal Organoid), hMO (Human Midbrain Organoid), hMG (Human Microglia-like Cells). (**B-I**) UMAP plots showing the expression of key marker genes across cell populations in different organoid models. (**B**) MAP2: Pan-neuronal marker, enriched in differentiated neurons. (**C**) DLX1: GABAergic interneuron marker, indicative of striatal and inhibitory neuronal identity. (**D**) SHH: Sonic hedgehog signaling, important for midbrain and ventral forebrain patterning. (**E**) SLC17A6 (VGLUT2): Excitatory glutamatergic neuron marker. (**F**) CAMK2A: Postsynaptic marker of excitatory cortical projection neurons. (**G**) GRIA2: AMPA receptor subunit, indicating excitatory neurotransmission. (**H**) GAD1: Enzyme involved in GABA synthesis, marking inhibitory GABAergic neurons. (**I**) GAD2: Another GABA synthesis enzyme, marking inhibitory interneurons.

**s-Figure 2.**
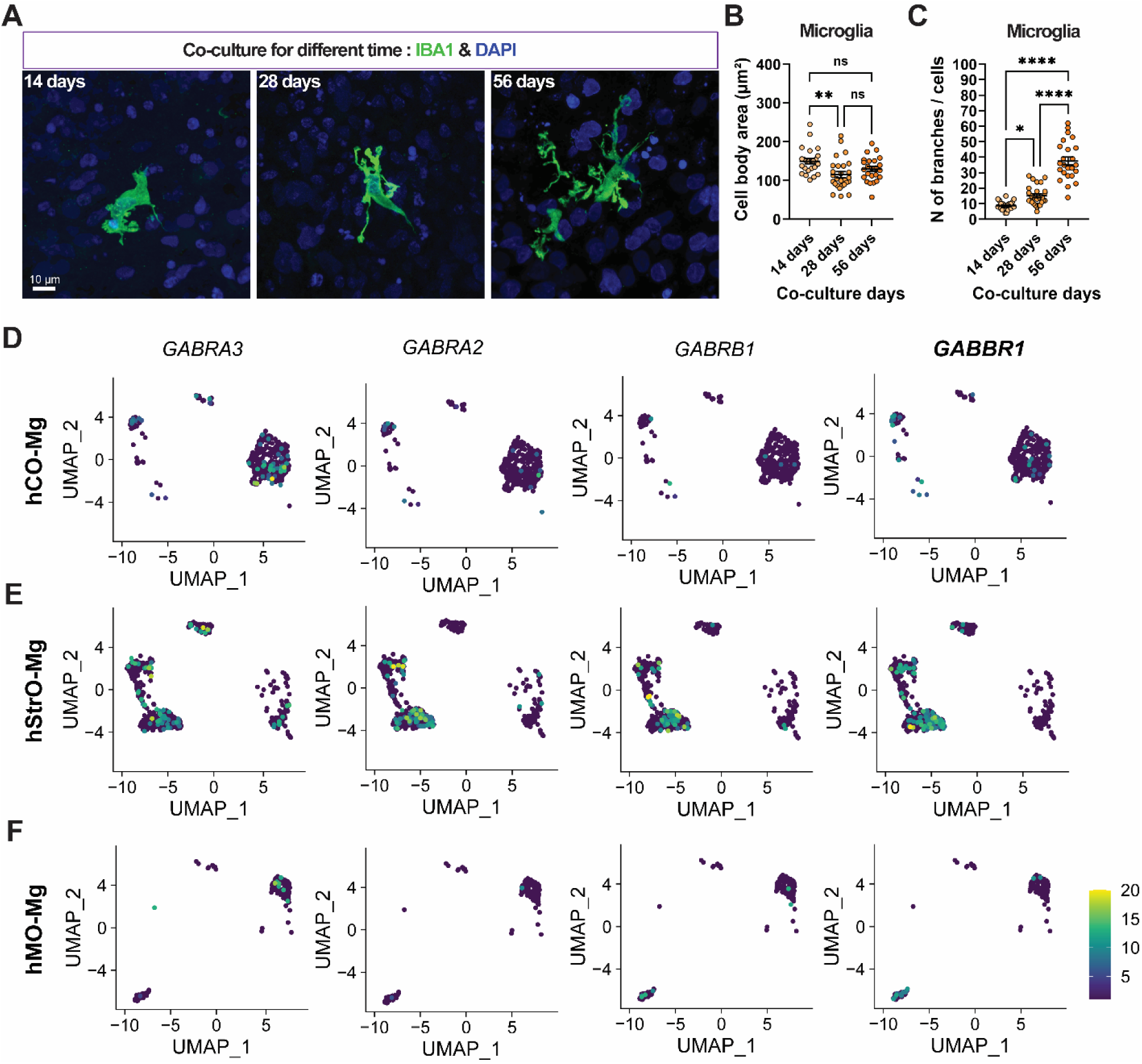
The maturation of striatal microglia increased with prolonged co-culture time, and the distributions of microglial GABA-related receptor genes. (**A**) Representative microglial morphology with increasing co-culture time in the hStrO-Mg group. Scale bar: 10 μm. (**B** and **C**) Quantifications of microglial cell body area and N of branches from each group. Co-culture time, 14 days group: n=24 cells/12 organoids, 28 days group: n=27 cells/12 organoids; 56 days group: n=24 cells/12 organoids. One-way ANOVA, *p* < 0.05 (*), *p* < 0.01 (**), *p* < 0.001 (***), and *p* < 0.0001 (****), ns: no significance. (**D**) The expression and distribution of GABA receptor-related genes in microglia UMAP from hCO-Mg scRNA-seq analysis. (**E**) The expression and distribution of GABA receptor-related genes in microglia UMAP from hStrO-Mg scRNA-seq analysis. (**F**) The expression and distribution of GABA receptor-related genes in microglia UMAP from hMO-Mg scRNA-seq analysis.

**s-Figure 3.**
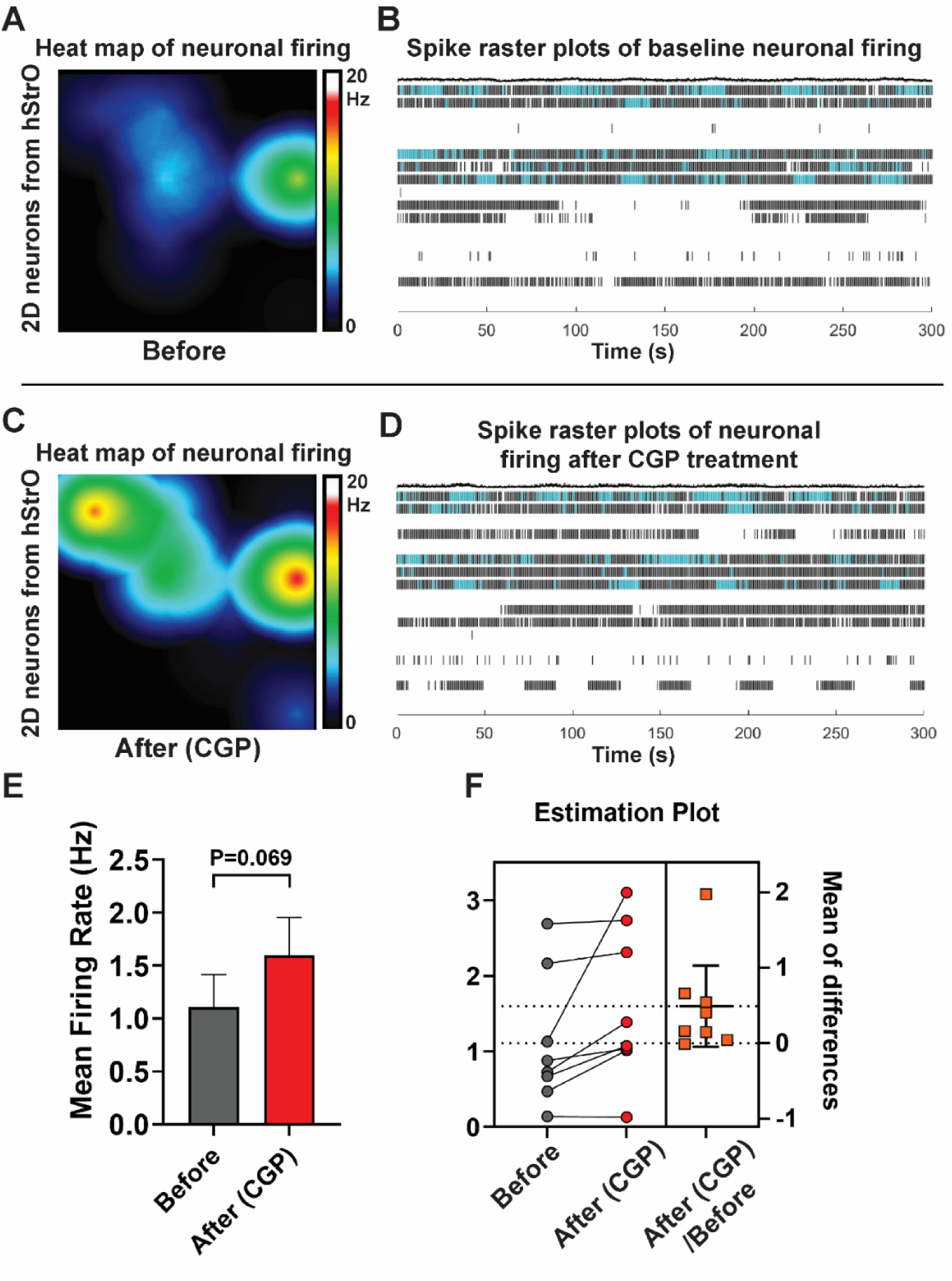
CGP treatment showed only a modest influence on neuronal excitability. (**A** and **C**) Representative heat map of neuronal firing including before (**A**) and after (**C**) CGP treatment. (**B** and **D**) Representative spike raster plots including before (**B**) and after (**D**) CGP treatment. Each row corresponds to the neuronal spike recordings obtained from a single electrode over a 300-second period, where each tick mark signifies a spontaneous event. Bursting events are represented by clusters of ticks highlighted in blue. (**E**) Quantifications of Mean firing frequency (MFF) comparison between before and after CGP treatment. n=8 wells. Paired Student’s t-test, *p* = 0.069. (**F**) Left: Neuronal mean firing frequency by MEA recording including before and after CGP treatment. Right: Within-subject differences, with the mean of difference represented by the solid horizontal line.

**s-Figure 4.**
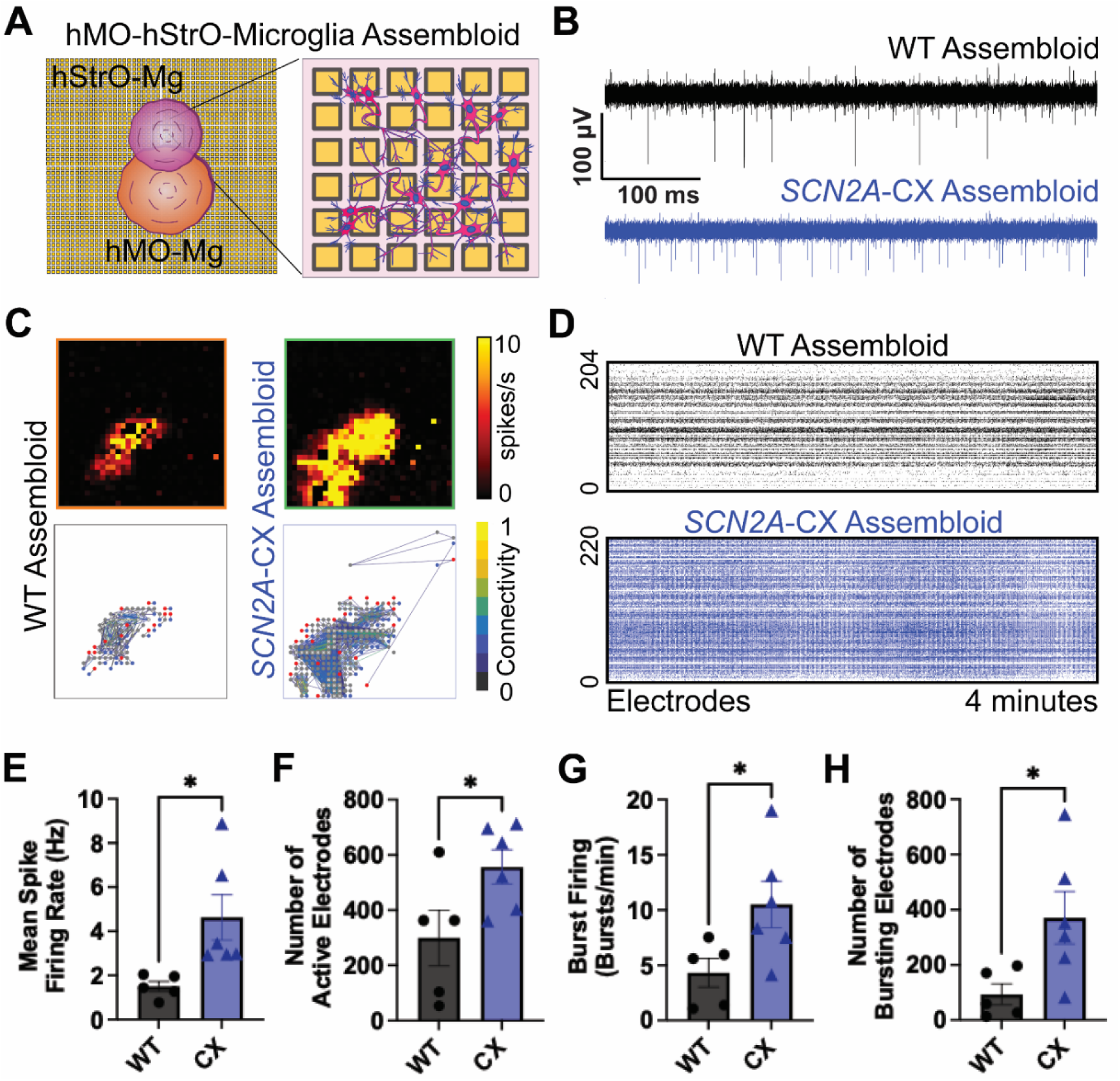
*SCN2A*-C959X renders network hyperexcitability in hMO-hStrO-Microglia assembloids. (**A**) Schematic of the co-culture system illustrating midbrain (orange) and striatal (pink) organoids plated onto a high-density microelectrode array (inset shows a zoomed view of individual electrodes and interconnections). (**B**) Example raw voltage traces from WT group (black) and *SCN2A* C959X-mutant group (blue) recorded by individual electrodes; note the increased spiking frequency in *SCN2A* C959X-mutant group. (**C**) Representative heat map of spike activity (spikes/s) (above) and representative neuronal connection map (below) by HD-MEA recording in WT group and *SCN2A* C959X-mutant group. (**D**) Spike raster plot compiled across ∼424 electrodes in WT group (black) and *SCN2A* C959X-mutant group (blue) networks over 4 minutes. Each row represents an electrode; each tick marks a detected spike. (**E**) Quantification of the mean spike firing rate (Hz). (**F**) Number of “active” electrodes in WT group vs. *SCN2A* C959X-mutant group. (**G**) Burst firing rate (bursts/min). (**H**) Number of “bursting” electrodes, defined as electrodes displaying bursting activity. E-H, WT group n=5 assembloids, CX group n=6 assembloids. Unpaired t-test, bars show mean ± SEM; *p* < 0.05 (*).

## Methods

### Generation of organoids (hCO, hStrO, and hMO) from hiPS cells

*SCN2A* c.2877C>A (p.Cys959Ter) mutant hiPSC lines were generated using CRISPR/Cas9- mediated genome editing(*50, 51*) in early passage (p2) KOLF2.1J reference iPSCs(*52*). We conducted recharacterization of pluripotency assays and genome integrity to ensure the quality of hiPSC(*53*). In total, five hiPSC lines were utilized: three isogenic control lines (KOLF 2.1J, B07 WT, and B11 WT) and two heterozygous *SCN2A*-C959X lines (A02 HET, E04 HET). The hiPSC colonies were maintained daily in StemFlex Medium (Gibco, A3349401) and then aggregated in Essential 8 medium (Gibco, A1517001) at a density of 100 cells/μL to form spheroidal embryoid bodies (EBs). Formation of the EBs was facilitated overnight in round-bottom ultra-low-attachment plates (Corning Costar, CLS7007) with a 3-minute centrifugation at 100 g. The EBs were subsequently cultured in Essential 6 medium (Gibco, A1516401) for the first 6 days.

To generate hCO, following the method described, EBs were initially maintained in E6 medium for 6 days with the addition of inhibitors targeting the activin/nodal/TGF-β and BMP pathways: dorsomorphin (2.5 μM, Sigma-Aldrich, P5499) and SB-4321542 (10 μM, R&D Systems, 1614)) as well as XAV-939 (1.25 μM, Tocris, 3748) to induce neuronal differentiation via the DUAL-SMAD approach. The EBs were then collected and transferred to flat-bottom six-well suspension culture plates (Corning, 3471) and maintained in Neurobasal-A medium (Thermo Fisher Scientific, 10888022), supplemented with Glutamax (Gibco, 35050061), Penstrep (10,000 U/mL, Gibco, 15140163), and B27 minus vitamin A (Gibco, 12587010), along with 20 μg/mL human recombinant FGF2 (R&D Systems, 233-FB-500) and 20 μg/mL EGF (R&D Systems, 236-EG). The medium was completely refreshed daily for the first 17 days, then every two days until day 23. Subsequently, the supplements were replaced with a formulation containing 20 ng/mL BDNF (PeproTech, 450-02), 20 ng/mL NT3 (PeproTech, 450-03), 50 μM cAMP (Santa Cruz, sc-201567A), 10 μM DHA (cis-4,7,10,13,16,19-docosahexaenoic acid) (MilliporeSigma, D2534), and 200 μM AA (L-ascorbic acid 2-phosphate trisodium salt) (Wako, 323-44822), with medium changes every two days until day 46. Afterward, the organoids were maintained in Neurobasal-A basal medium with B27+ supplement (without additional growth factors) until day 150, with media changes every 4-5 days.

For the generation of hStrO, the protocol previously outlined was followed(*54*). On day 6, the EBs were transferred into a neural medium composed of Neurobasal-A, B-27 without vitamin A, GlutaMAX, and penicillin-streptomycin. This medium was also supplemented with 2.5 μM IWP-2 (a WNT pathway inhibitor; Selleck Chemicals, S7085) and 50 ng/mL recombinant Activin A (PeproTech, 120-14P). On day 11, the retinoid X receptor agonist SR11237 (100 nM, Tocris, 3411) was added alongside the existing supplements. Starting from day 22, to promote differentiation of neural progenitors into neurons, the medium was further enriched with 20 ng/mL BDNF, 20 ng/mL NT-3, 200 μM AA, 50 μM cAMP, and 10 μM DHA. From day 42 onward, DAPT (2.5 μM, Stemcell Technologies, 72082) was added in conjunction with BDNF, NT-3, AA, cAMP, and DHA. From day 46, the cultures were maintained solely in a neural medium containing B-27 Plus Supplement with medium changes every 4 days.

To generate hMO(*55*), EBs were cultured in brain organoid generation medium (BGM) containing a 1:1 mix of DMEM/F12 (Fisher, 21331020) and Neurobasal A Medium, 1% penicillin/streptomycin (PS), 1% GlutaMAX, 1% MEM Non-Essential Amino Acids Solution (NEAA) (Thermo Fisher Scientific, 11140050), 55 μM β-mercaptoethanol (Fisher, 21985023) and 1 μg/ml heparin (Sigma, H3149). The medium was supplemented with 1% N-2 Supplement (Fisher, 17502048) and 2% B27 without vitamin A supplement, 2 μM dorsomorphin, A83-01 (2 μM, PeproTech, 9094360), CHIR99021 (3 μM, Tocris, 4423), and IWP2 (1 μM, Selleck Chemicals, S7085) for 7 days. On day 4, 100 mg/ml FGF8b (PeproTech, AF-100-25) and Smoothened Agonist (SAG) (2 μM, PeproTech, 9128694) were added, and on day 7, 200 ng/ml Laminin (Sigma, L4544) and 2.5 μg/ml insulin (Sigma, I9278) were introduced, with these supplements maintained until day 9. After this period, the medium was switched to BGM containing 1 % N-2, 2% B27 plus, 10 ng/mL BDNF, 10 ng/mL GDNF (PeproTech, 450-10), 125 μM cAMP, and 200 μM AA. The cultures were maintained under these conditions for up to 6 months.

### Human microglia differentiation

Microglia differentiation was performed based on a previously established protocol(*56*). In summary, iPSCs (Kolf2.1J endogenously expressing GCaMP6f, B11 WT, and B07 WT) were first differentiated into hematopoietic progenitor cells over 12 days using the STEMdiff Hematopoietic Kit (STEMCELL TECHNOLOGIES, 05310). iPSCs were grown in a 6-well plate coated with Matrigel (Corning 354277) and maintained in mTeSR™ Plus medium (STEMCELL TECHNOLOGIES, 100-0276). The cells were passaged by washing with 1x PBS followed by the addition of 1 mL of ReLeSR (STEMCELL TECHNOLOGIES, 100-0483) and incubated at 37°C for 90 seconds. ReLeSR was then removed and 1 mL of mTeSR Plus was pipetted over the cells. 30-50 colonies were plated on one well of a Matrigel-coated 6-well plate using a wide-bore pipette tip. One day after cell seeding Hematopoietic progenitor medium A was added to the cells. After 2 days a half medium change was done. On day 3 medium A was removed, and Hematopoietic progenitor medium B was added. A half-medium change was done on days 5, 7, and 10. On day 12 the progenitor cells were harvested from the well by pipetting several times before transfer to one well of a Matrigel coated 6 well plate. The resulting progenitor cells were then transferred into 2 mL of microglia differentiation medium composed of DMEM/F12 (Gibco, 11320033), 2× insulin-transferrin-selenite (Gibco, 41400045), 2× B27 plus (Gibco, A3582801), 0.5× N2 (Gibco, 17502048), 1× non-essential amino acids (Gibco, 11140050), 400 μM monothioglycerol (Sigma, M6145-25MG), 5 μg/mL human insulin (Sigma I2643-25MG). Prior to use, the differentiation medium was supplemented with 100 ng/mL IL-34 (Gibco, 200-34-10UG), 50 ng/mL TGF-β1 (100-21-10UG), and 25 ng/mL M-CSF (Gibco, 300-25-10UG). The cells were maintained in this medium for up to 24 days with the addition of 1 mL of media every 2 days.

### Microglia integration in the cortical, striatum, and midbrain organoids

To facilitate the incorporation of microglia into the organoids (hCO, hStrO, and hMO), organoids that had cultured for over 80 days were individually transferred to an ultra-low attachment 96- well plate. On day 12 of microglial differentiation, microglia were collected and seeded at a density of 400,000 cells per well onto the surface of the organoids in fresh medium composed of an equal mixture of microglial differentiation medium and mature cortical/striatal/midbrain organoid medium. The organoids were incubated for 7 days to allow spontaneous microglial integration, with daily medium replacement. Afterward, the organoids were moved back to an ultra-low attachment six-well plate and maintained for an additional 7 days following the standard cortical/striatal/midbrain organoid culture protocol.

### Generation of microglial integrated assembloids

To produce either hMO-hStrO-Microglia or hCO-hStrO-Microglia assembloids, we first generated hMO-Mg (or hCO-Mg) and hStrO-Mg separately from hiPSCs. Next, these microglia-containing organoids were combined by placing them in close proximity within 1.5LmL Eppendorf tubes and incubating them for 7 days. During this incubation period, half of the medium was gently replaced each day. On the seventh day, the assembloids were transferred using a trimmed P1000 pipette tip into 24-well ultra-low attachment plates, and the medium was subsequently refreshed every 2 days.

### Viral labeling and live cell imaging

Prior to assembling the two distinct microglia-integrated organoids, 3D neural organoids were virally transduced to both visualize projections and trigger neuronal hyperexcitability using pAAV1-hSyn-mScarlet (Addgene, 131001) and pAAV9-hSyn-hM3(Gq)-mCherry (Addgene, 50474), respectively. For the hMO-hStrO-Microglia assembloid experiments, hMO-Mg were first transferred to a 24-well plate containing 200 μl of medium and incubated overnight with the virus in an incubator. The following day, 800 μl of fresh culture medium was added. Three days post-infection, the virus-labeled hMO-Mg and hStrO-Mg were utilized for assembloid formation. Live cell imaging was performed on the midbrain-striatal projection area at assembly days 10, 20, and 30 using a confocal fluorescence microscope equipped with an incubation system (LSM900; Carl Zeiss, Jena, Germany). Image data were subsequently analyzed with Fiji (ImageJ).

### Chemogenetics stimulation and calcium imaging

For chemogenetics experiments, hMO-hStrO-Microglia assembloids—with hMO previously transduced with pAAV-hSyn-hM3(Gq)-mCherry—were placed on a 20Lmm coverslip (within a 35Lmm glass-bottom plate) in neural medium and imaged using a 10× objective on the LSM900 Zeiss confocal microscope. To induce neuronal hyperexcitability specifically in the hMO region, 10LμM clozapine N-oxide (targeting the Gq-DREADD) (Abcam, ab120019), was applied. Calcium activity, monitored via GCaMP6f, was recorded for a total duration of 5 minutes and 30 seconds; baseline activity was captured for the initial 100 seconds, after which CNO was added, and recording continued for an additional 230 seconds. To inhibit microglial GABA_B_ receptor signaling, the medium was supplemented with 2 μM CGP 55845 hydrochloride (a selective GABA_B_ receptor antagonist) (MCE, HY-103516), with the medium refreshed every 6 hours prior to calcium imaging.

### Calcium activity analysis

Calcium imaging data was recorded from regions of interest (ROIs) in assembloids samples and exported as Excel files. To define ROIs for microglial somas and processes, the user manually delineated these areas on an average intensity projection covering the full field of view using Inscopix. Next, the somatic ROIs were masked in the average intensity image to separate them from the microglial processes. By applying a threshold to the remaining image data, process ROIs were isolated, and their mean pixel intensities were quantified in a similar manner. Each column in the dataset represents the fluorescence intensity of a single soma or process over time. For quantifying calcium transients, fluorescence intensity values were converted into ΔF/F values as: ΔF/F=(F(t)−F0)/F0. For each soma and process, a 100-second moving window was applied to create a dynamic bassline. The baseline fluorescence (F0) is established by the 25th percentile of the intensity values within the moving window. Calcium transients were identified using a threshold-based method, and frames exceeding a threshold that is three times greater than the standard deviation of the baseline were classified as active transients. Maximum amplitude was computed by the peak ΔF/F value during detected transients. Signal area is the cumulative ΔF/F value above the threshold across all detected transients.

### Single-cell RNA-seq library preparation and data analysis

Organoids were dissociated into single-cell suspensions following standard protocols with modifications optimized for the PIPseq-T20 workflow. Briefly, 4-5 randomly selected organoids were pooled and incubated in an enzymatic dissociation solution containing 30 U/mL papain (Worthington Biochemical, LS003126) and 0.4% DNase I (Worthington Biochemical, LS2007) at 37°C for 45 min. Following enzymatic digestion, organoids were washed in protease inhibitor-containing medium and gently triturated to achieve a single-cell suspension. The cell suspension was filtered through a 70 μm Flowmi Cell Strainer (Bel-Art, H13680-0070) and counted. Viable cells were resuspended in Cell Suspension Buffer (Fluent BioSciences, PIPseq T20 v3.0) at a final concentration of 10000 cells/μL, ensuring optimal loading efficiency. Single-cell RNA-seq libraries were generated using the PIPseq-T20 3L Single Cell RNA Kit v3.0 (Fluent BioSciences) following the manufacturer’s protocol. Briefly, 40,000 cells per reaction were loaded into PIPseq Pre-templated Instant Partitions (PIPs) and mixed with partitioning reagents to encapsulate individual cells with barcoded beads. Cell lysis and mRNA capture were performed within the PIPs, followed by cDNA synthesis using a template-switch oligonucleotide (TSO) approach. cDNA was then amplified using whole transcriptome amplification (WTA) and purified via SPRI bead cleanup. Purified cDNA was fragmented, end-repaired, and A-tailed prior to adapter ligation using the PIPseq-T20 Library Preparation Kit. Libraries were indexed using dual-index P7/P5 adapters, amplified via PCR, and size-selected using double-sided SPRI bead purification. The final libraries were quantified using a Qubit High Sensitivity DNA Assay Kit (Thermo Fisher, Q33231) and analyzed on a Bioanalyzer 2100 or TapeStation 4200 (Agilent). Sequencing was performed on an Illumina NovaSeq S4 platform with 2 × 150 bp paired-end reads, targeting ∼20,000 reads per cell.

The resulting feature-barcode matrices were read into R (version 4.2.2), excluding any cell expressing fewer than 200 genes and any gene expressed in fewer than three cells. For all single-cell samples, cells with greater than 15% mitochondrial, fewer than 2,500 features or more than 10,000 features were removed by Seurat (version 4.3.0.1). Similarly, cells with fewer than 2,000 or more than 50,000 counts were filtered out. We merged the samples with the “merge” function. Then, total cell clustering was performed by the “FindNeighbors” and “FindClusters” functions using the first 50 PCs and a resolution of 0.2, and for visualization with UMAP. Clusters were grouped based on the expression of known marker genes. The classification of microglia was performed based on the combinatorial expression of known markers as previously described(*39*).

### High Density Multielectrode Arrays (HD-MEA) recording

Multiwell HyperCam Alpha (3Brain AG, Switzerland) High Density Multielectrode Arrays were used for all HD-MEA recordings. The 6-well plates were cleaned using 200uL 1% Tergazyme solution for 1 hour at 37°C, washed 3 times with excess Sterile MilliQ water (R > 18.2), then disinfected with 70% ethanol for 1 hour. Ethanol was removed and plates were dried under UV, then the plates were incubated overnight in PBS at room temperature. Next, HD-MEAs were coated sequentially with poly-L-ornithine (50 μg/mL) in sterile MilliQ overnight, washed 3 times with MilliQ water, and then incubated with Laminin (50 μg/mL) for 4 h at 37°C. Day 100-150 Assembloids were then seeded onto the 3Brain MEAs in 20 μL media and returned to the incubator for 2 hours. An additional drop of 20 μL media was added using wide bore pipette tips every 2 hours for 8 hours. Finally, 2 mL of media was gently added in concentric circles to fill the well. Media was exchanged every 3-4 days. Assembloids were recorded between days 10-15. Data was acquired and analyzed using Brainwave V software, (v5.6, 3Brain Switzerland). HD-MEA arrays were recorded using a 2304 × 2304 electrode configuration, with a 60 μm pitch (2.9 × 2.9 mm^2^ area). The sampling rate was set to 10,000 Hz, and a hardware high-pass filter of 100Hz was used. Direct light stimulation compensation was applied during the recording. For Spike Detection Band-pass Filter of 20-5000Hz, using Fast Fourier Transform (FFT) was performed. Spike detection was set to a Standard Deviation of 8.0 with a Peak Lifetime of 2.0 ms, and a Refractory Period of 1.0 ms. Pre-Peak Wave Duration was 1.0 ms. Electrodes with less than 0.083 Hz (5 spikes per minute) were discarded. For spike burst detection, simple interspike interval (ISI) settings were used with a max ISI of 100 ms and the minimum number of spikes set to 5. Spike Sorting using 3 component PCA with K-Means & Gap Statistics clustering was applied. For spike network bursts, a recruitment-based algorithm was applied with 10% recruited electrodes and a minimum spike threshold of 50 spikes. A spike cross-correlation window of 30 ms, and bin size of 3.0 ms was used to generate correlation matrices.

### Cryosection and immunofluorescence

Organoids and assembloids were initially fixed in a solution of 4% paraformaldehyde (PFA) in phosphate-buffered saline (PBS) for 4 hours. After fixation, the samples were transferred into a 30% sucrose/PBS solution and left for 1 day until they sank to the bottom. Once dehydration was complete, the samples were embedded in an OCT/30% sucrose/PBS mixture and stored at -80°C for subsequent experiments. For immunofluorescence staining, 30Lμm-thick sections were prepared using a Leica Cryostat (Leica, CM1860). The cryosections were first rinsed with PBS to remove any residual OCT and then blocked for 1 hour at room temperature in a solution containing 10% Normal Donkey Serum (NDS; Millipore Sigma, S30-100ML), 0.3% Triton X-100 (Millipore Sigma, T9284-100ML), and 1% BSA diluted in PBS. Next, the sections were incubated overnight at 4°C with the appropriate primary antibody solution. On the following day, the sections were washed with 0.01 M PBS and then incubated with secondary antibodies for 2 hours at room temperature. After a final wash, coverslips were mounted onto the sections. The primary antibodies used were as follows: anti-IBA1 (Abcam, ab178846), human anti-IBA1 (Synaptic Systems, 234308), human anti-CD68 (Invitrogen, MA5-13324), human anti-Gabbr1 (Gene Tex, GTX102511), human anti-DRD1 (Thermo Fisher, 702593), anti-Ctip2 (Abcam, ab18465), anti-TBR1 (Thermo Fisher, 66564), anti-GABA (Thermo Fisher, PA5-32241), anti-TH (Thermo Fisher, MA1-24654), anti-MAP2 (Thermo Fisher, PA1-10005), anti-NeuN (Thermo Fisher, 702022), anti-GAD67 (Thermo Fisher, PA5-21397), anti-FOXA2 (Thermo Fisher, 701698), and anti-OTX2 (Thermo Fisher, MA5-15854). Confocal images were acquired using a Z-stack laser scanning confocal fluorescence microscope (LSM900; Carl Zeiss, Jena, Germany). Three-dimensional image analysis was conducted with Imaris 9.9 software. The reconstructed surfaces of IBA1 and CD68 were measured, and the percentage of CD68 occupancy within microglia was determined using the formula: (volume of CD68) / (volume of IBA1^+^ cell). Additionally, to quantify the VGAT^+^ signal within CD68^+^ regions, the following calculation was used: (VGAT^+^ volume / CD68^+^ volume) × 100.

### Electrophysiology

Patch-clamp recordings were performed as described previously(*32, 35*). Briefly, organoids were cut with a vibratome (Leica VT1200S, Germany) in ice-cold slicing solution containing (in mM): 110 choline chloride, 2.5 KCl, 1.25 NaH_2_PO_4_, 25 NaHCO_3_, 0.5 CaCl_2_, 7 MgCl_2_, 25 glucose, 1 sodium ascorbate and 3.1 sodium pyruvate (pH 7.4, 305-315 mOsm, bubbled with 95% O_2_ and 5% CO_2_). Slices were incubated in the same medium for 10 minutes at 33°C, then transferred to artificial cerebrospinal fluid (aCSF; in mM; 125 NaCl, 2.5 KCl, 1.25 NaH_2_PO_4_, 25 NaHCO_3_, 2.0 CaCl_2_, 2.0 MgCl_2_, 10 glucose; pH 7.4, 305-315 mOsm, bubbled with 95% O_2_ and 5% CO_2_) for 10-20 minutes at 33°C before storage at room temperature for at least 30 minutes. Slices were visualized under IR-DIC using a BX-51WI microscope (Olympus) with an IR-2000 camera (Dage-MTI). We used thin-wall borosilicate pipettes (BF150-110-10) with 3-5 MΩ open-tip resistances. For current-clamp recordings, the internal solution contained (in mM): 122 KMeSO_4_, 4 KCl, 2 MgCl_2_, 0.2 EGTA, 10 HEPES, 4 Na_2_ATP, 0.3 Tris-GTP, 14 Tris-phosphocreatine, adjusted to pH 7.25 with KOH, 295-305 mOsm. Recordings were performed with an Axon MultiClamp 700B amplifier (Molecular Devices), and data were acquired using pClamp 11.3 software filtered at 2 kHz and sampling rate at 50kHz with an Axon Digidata 1550B plus HumSilencer digitizer (Molecular Devices). The action potentials were obtained in response to a series of 400-ms hyperpolarizing and depolarizing current steps from -200 pA to +400 pA in increments of 50-pA, each sweep duration of 5 s with cells held at the normal RMP.

### Statistical analysis

OriginPro 2025 and GraphPad Prism 10 were used for data analysis and curve fitting. Two-tailed Student’s t-test (parametric) or unpaired two-tailed Mann-Whitney U-test (non-parametric) was used for single comparisons between two groups. The other data were analyzed using one-way or two-way ANOVA and then using a post hoc with Bonferroni corrections. The number of experimental samples (n) in each group was indicated in the legend. Results are presented as mean ± standard error of the mean (SEM). Significance was determined when *p* < 0.05 (*), *p* < 0.01 (**), *p* < 0.001 (***), and *p* < 0.0001 (****).

